# SCREEN: a graph-based contrastive learning tool to infer catalytic residues and assess mutation tolerance in enzymes

**DOI:** 10.1101/2024.06.27.601004

**Authors:** Tong Pan, Yue Bi, Xiaoyu Wang, Ying Zhang, Geoffrey I. Webb, Robin B. Gasser, Lukasz Kurgan, Jiangning Song

**Affiliations:** Biomedicine Discovery Institute and Department of Biochemistry and Molecular Biology, Monash University, Melbourne, Australia; Department of Data Science and Artificial Intelligence, Monash University, VIC 3800, Australia; School of Computer Science and Engineering, Nanjing University of Science and Technology, Nanjing, China; Department of Veterinary Biosciences, Melbourne Veterinary School, The University of Melbourne, Parkville, Australia; Department of Computer Science, Virginia Commonwealth University, Richmond, USA

**Keywords:** catalytic residue, enzyme structure, evolutionary conservation, graph neural network, contrastive learning

## Abstract

The accurate identification of catalytic residues contributes to our understanding of enzyme functions in biological processes and pathways. The increasing number of protein sequences necessitates computational tools for the automated prediction of catalytic residues in enzymes. Here, we introduce SCREEN, a graph neural network for the high-throughput prediction of catalytic residues via the integration of enzyme functional and structural information. SCREEN constructs residue representations based on spatial arrangements and incorporates enzyme function priors into such representations through contrastive learning. We demonstrate that SCREEN (i) consistently outperforms currently-available predictors; (ii) provides accurate results when applied to inferred enzyme structures; and (iii) generalizes well to enzymes dissimilar from those in the training set. We also show that the putative catalytic residues predicted by SCREEN mimic key structural and biophysical characteristics of native catalytic residues. Moreover, using experimental data sets, we show that SCREEN’s predictions can be used to distinguish residues with a high mutation tolerance from those likely to cause functional loss when mutated, indicating that this tool might be used to infer disease-associated mutations.

## Introduction

Enzymes are critical for a wide range of diverse biochemical, molecular and physiological processes and pathways which sustain life (*1*). The extraordinary catalytic proficiency of an enzyme is often intricately orchestrated by a selected set of amino acids within its active site(s), referred to as the catalytic residues (*2*). These spatially proximate catalytic residues can engage in precise, non-covalent interactions with substrate molecules, catalyzing chemical reactions and ensuring the catalytic efficiency and specificity of enzymes (*3*). Catalytic residues often exhibit conservation across species, particularly those within the same taxonomic groups (*4*), such that mutations in catalytic sites can affect enzyme function(s), potentially inducing the onset of diseases, such as cancers and metabolic disorders (*5*). For instance, mutations of catalytic residues in the tumor suppressor phosphatase and tensin homolog (PTEN) have been shown to culminate in various forms of cancers, such as glioblastoma multiforme, melanoma and breast cancer (*6*). Mutations in the catalytic sites of enzymes, such as CYP2C9, which are responsible for the biotransformation of small molecule drugs, can impact individual drug responses and potentially increase the risk of metabolic disorders (*7*).

The number for which enzymes with detailed catalytic residue annotations available in the Mechanism and Catalytic Site Atlas (M-CSA) database (*8*) is substantially lower than the vast number of enzyme sequences in the Swiss UniProt database (∼64,815 entries) and enzyme structures in repositories such as the Protein Data Bank (PDB) (∼25,868 entries with assigned EC numbers) (*9, 10*). This gap relates primarily to the absence of high-throughput methods for identifying catalytic residues. Traditionally, the active sites in enzymes have been established using site-directed mutagenesis and biochemical assays, providing insight into the corresponding kinetic and thermodynamic parameters (*11*). However, these laboratory methods are relatively low throughput, time-consuming and labor-intensive, thereby restricting analyses to small numbers of residues and constraining ‘down-stream’ applications such as the design of novel enzymes and inhibitors (*12*). Furthermore, although M-CSA offers information on enzyme functions, its coverage of the function space is not comprehensive, particularly for oxidoreductases and translocases (with only 45.6% and 30% coverage, respectively), potentially attributable to curation backlog and/or limited functional data/information (*13*). There is significant demand for an *in silico* approach for the reliable and reproducible identification of catalytic residues in enzymes from sequence and/or structure data to enable the exploration of enzyme functions and accelerate biomolecular design.

A significant effort has been directed towards developing computational methods for the identification of catalytic residues in enzymes (*14*). These methods utilize the ‘ground truth’ annotations of catalytic residues from curated public databases to train predictive models, which, in turn, can be applied to infer catalytic residues in most unknown enzymes (*15*). For one of the earliest tools, Gutteridge et al. (*16*) trained a neural network model to identify catalytic residues using enzyme structure- and sequence-derived features as inputs. Subsequently developed predictors often employed support vector machine (SVM) models or random forest models to a collection of manually curated features (*17-21*). Interestingly, Chea et al. (*22*) did not use a machine learning model but, instead, predicted catalytic residues using statistical scores calculated from a network representation of protein structure and solvent accessibility. However, the models often did not explicitly consider the spatial organization of amino acid residues or fully investigate the relationship between catalytic residues, enzyme structure and enzyme function, thus limiting the predictive capacity of these computational tools.

We anticipate that recognizing patterns in the spatial arrangement of residues within enzyme structures can substantially enhance the performance of catalytic residue prediction tools and deepen our understanding of enzyme function through the use of modern deep neural networks. To this end, we propose here a deep learning-based solution, called SCREEN, for the accurate prediction of catalytic residues in enzymes. SCREEN employs a graph neural network that models the spatial arrangement of active sites in enzyme structures and combines data derived from enzyme structure, sequence embedding and evolutionary information obtained by using two complementary methods – BLAST (Basic Local Alignment Search Tool) and HMMER (sequence analysis tool using profile hidden Markov models) (*23, 24*). Moreover, we apply the contrastive learning framework to further enhance the predictive performance of SCREEN by incorporating enzyme functional information.

## Results

### SCREEN model

First, we curated a large dataset of 1055 enzymes from several public databases and literature. This dataset included protein structure and sequence data, which we used as inputs for the model, as well as annotations of catalytic residues and Enzyme Commission (EC) numbers (*25*), which we utilized to train the model. SCREEN is a supervised deep learner that integrates information derived from atomic structures, sequences and evolutionary profiles (Fig. 1). Specifically, SCREEN presents the input (enzyme structures) as graphs at the residue level, leveraging evolutionary information, sequence embeddings – generated by a modern language model – and relevant structural characteristics, such as B-factors and solvent accessibility (*26*). We correspondingly employed a graph convolutional neural network to generate propensities for catalytic residues from these inputs. The training process employed enzyme function information (EC main class) via a contrastive learning framework, utilizing the Triplet Margin Loss function to enable clustering enzymes of the same classes and separating enzymes from different classes in a latent feature space. This allowed our model to develop class-specific latent feature spaces, leading to improvements in predictive performance/capacity. Moreover, we employed a dynamic training strategy, in which we initially trained the network using the contrastive learning and then applied our model to accurately identify catalytic residues.

**Fig. 1.**
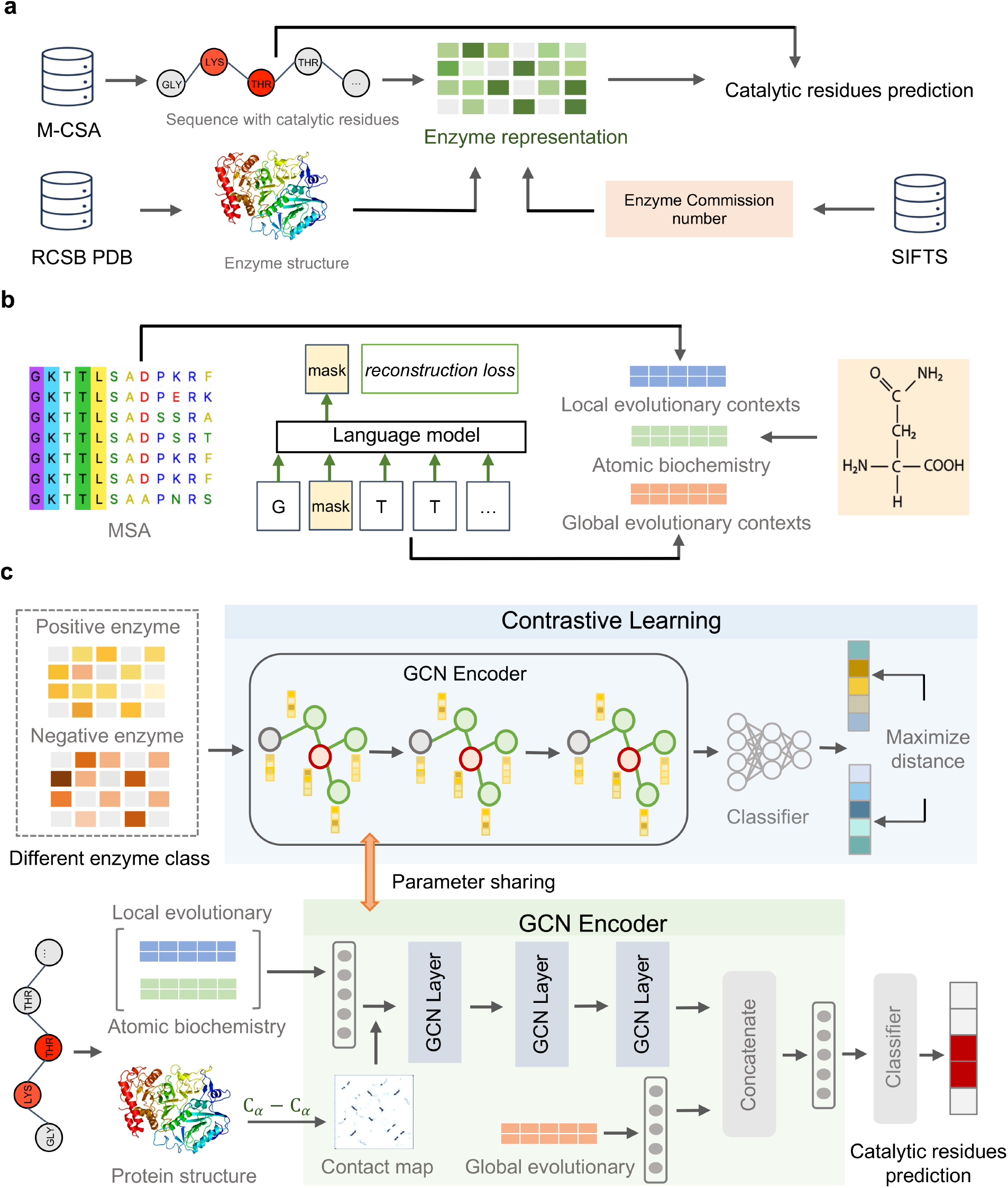
SCREEN – a predictor of catalytic residues in enzymes. (**A**) Data collection. We curate catalytic residue annotations from the M-CSA database, extract the corresponding enzyme structures from the RCSB PDB database, and retrieve corresponding enzyme function information using the SIFT program (*62*). (**B**) Generation of inputs that include evolutionary profiles based on multiple sequence alignments (MSA), sequence embeddings that leverage a large-scale protein language model, and structural characteristics derived from the atomic structures. (**C**) The architecture of SCREEN’s predictive model.

### SCREEN accurately predicts catalytic residues

We collected five commonly-used test datasets to comparatively assess SCREEN against seven current solutions. These test datasets included the EF superfamily and EF fold datasets (*20*), the HA superfamily dataset (*22*), the NN dataset (*16*) and the PC dataset (*18*). We compared the results from SCREEN with those of a conventional sequence-based method, CRpred (*17*), and six tools that employed different predictive models based on enzyme structures. These tools included a neural network-based approach (*16*), three SVM-based methods (*18*), a random forests-based PREvaIL (*21*), and a statistical approach (*22*). We calculated precision and recall as well as the F1 score to evaluate the predictions of catalytic residues (see Table 1). We showed that SCREEN consistently outperformed the seven tools for the five test datasets. Compared with the second-best method, SCREEN achieved an F1 score of 61.5 vs. a second-best F1 score of 32.8 for the EF superfamily dataset; 64.5 vs. 31.5 for the EF fold dataset; 72.0 vs. 45.0 for the HA superfamily dataset; 73.8 vs.36.5 for the NN dataset; and 74.1 vs. 26.3 for the PC dataset. The high F1 scores achieved by SCREEN were coupled with balanced and high values of precision and recall that ranged between 61.0 and 69.3 and between 61.2 and 82.0, respectively. We also quantified and compared two other popular metrics, the area under the receiver operating curve (AUC) and the area under the precision-recall curve (AUPR). Fig. 2a reveals that SCREEN achieved substantially higher AUC and AUPR scores when compared with the latest structure-based (PREvaIL) and the sequence-based (CRpred) tools, except for the EF superfamily and EF fold dataset, where AUC and AUPRC values were comparable.

**Table 1.** Comparison with current predictors of catalytic residues.

| Methods | Measureme<br>nt | EF<br>Superfamily<br>dataset | EF fold<br>dataset | HA<br>superfamil<br>y<br>dataset | NN<br>dataset | PC<br>dataset |
| --- | --- | --- | --- | --- | --- | --- |
| Methods from Youn et al. (20); Chea et al. (22); Gutteridge et al. (16); and Petrova and Wu (18) | Precision (Recall) | 16.9 (53.9)<sup>a</sup> | 17.1 (51.1)<sup>a</sup> | 16.5 (29.3)<sup>b</sup> | 56.0 (14.0)<sup>c</sup> | 7.0 (90.0)<sup>d</sup> |
|  | F1 | 25.7 | 25.6 | 21.1 | 22.4 | 13.0 |
| CRpred (17) | Precision (Recall) | 15.9 (52.1)<sup>e</sup> | 16.1 (48.0)<sup>e</sup> | 24.7 (49.7)<sup>e</sup> | 65.9 (18.0)<sup>e</sup> | 5.6 (84.5)<sup>e</sup> |
|  | F1 | 24.4 | 24.1 | 33 | 28.3 | 10.5 |
| CRHunter (19) | Precision (Recall) | 21.5 (68.7)<sup>f</sup> | 21.0 (62.7)<sup>f</sup> | 33.2 (69.6)<sup>f</sup> | 76.4 (24.0)<sup>f</sup> | 7.6 (92.1)<sup>f</sup> |
|  | F1 | 32.8 | 31.5 | 45.0 | 36.5 | 14.0 |
| PREvalL (21) | Precision (Recall) | 17.0 (59.4)<sup>g</sup> | 17.0 (56.5)<sup>g</sup> | 17.0 (57.9)<sup>g</sup> | 58.9 (17.0)<sup>g</sup> | 17.0 (58.1)<sup>g</sup> |
|  | F1 | 26.4 | 26.1 | 26.3 | 26.4 | 26.3 |
| SCREEN (This study) | Precision (Recall) | 61.9 (61.2) | 61.0 (68.4) | 69.3 (74.8) | 68.5 (79.9) | 67.6 (82.0) |
|  | F1 | 61.5 | 64.5 | 72.0 | 73.8 | 74.1 |

Results for: <sup>a</sup> EF superfamily and EF fold datasets (20); <sup>b</sup> HA family dataset (22); <sup>c</sup> NN dataset using the structure-based method without spatial clustering from Gutteridge et al. (16); <sup>d</sup> PC dataset from Petrova and Wu (18); <sup>e</sup> five test datasets from Zhang et al. (17); <sup>f</sup> five test datasets from Sun et al. (19); <sup>g</sup> five test datasets from Song et al. (21).

**Fig. 2.**
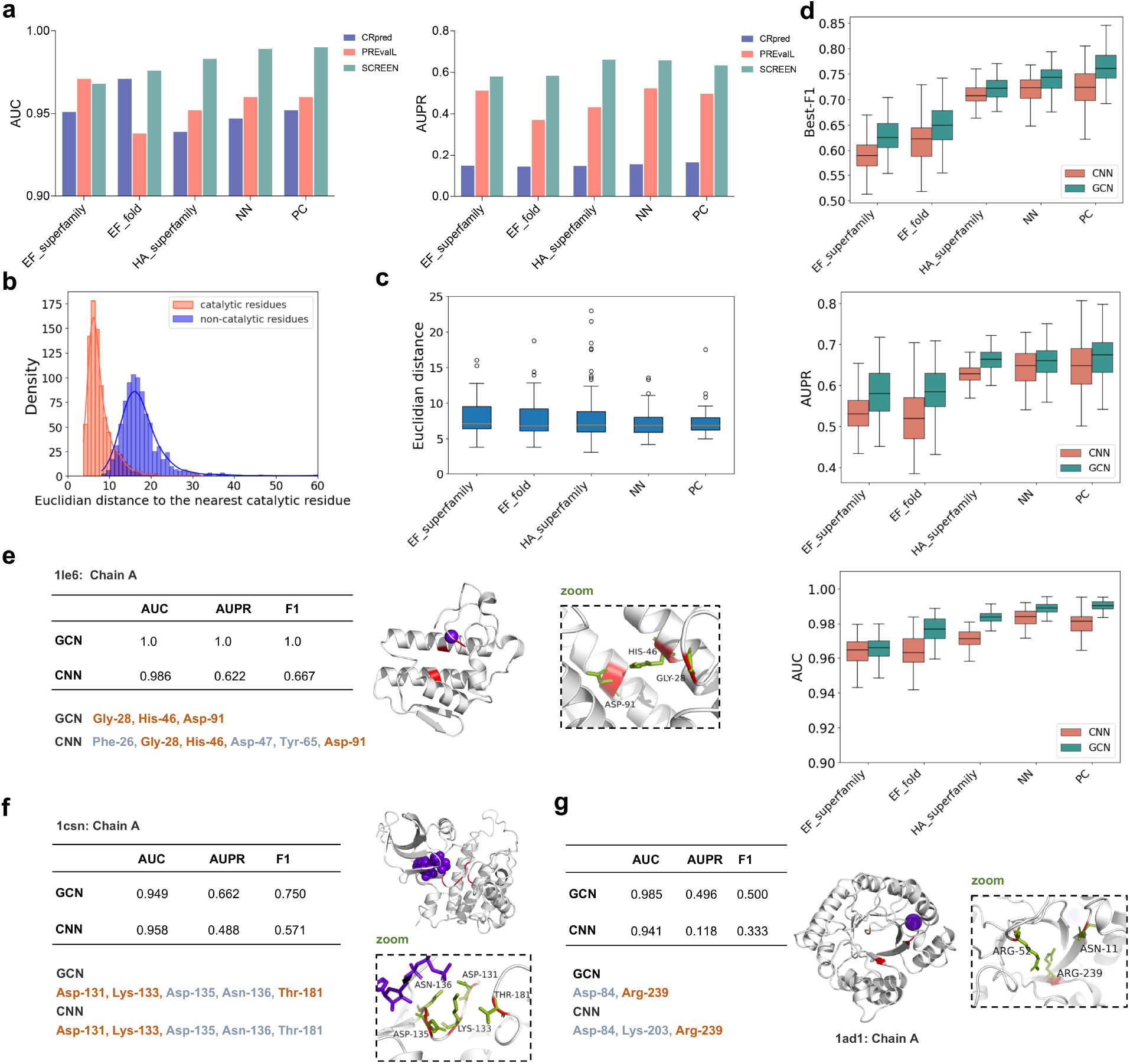
Predictive performance of SCREEN. (**A**) Comparison of the sequence-based CRpred and the newest structure-based PREvaIL with AUC and AUPRC metrics using each of five test datasets. **(B)** The distribution of the Euclidean distances to the nearest catalytic residues for the residues in the dataset of enzymes from the M-CSA database (*13*) (**C**) The distribution of the Euclidean distances to the nearest catalytic residue (using a boxplot) for the catalytic residues predicted by SCREEN. For boxplots, the center line represents the median, top and bottom edges are the first and third quartiles, respectively. **(D)** Comparison of the structure-based SCREEN model (GCN encoder) with a baseline model that excludes structure input and uses convolutional neural network (CNN encoder) based on the AUPRC, Best-F1 and MCC metrics on the five test datasets. **(E-G)** Three examples, ranked approximately at 10% (E), 50% (F), and 90% (G) based on the Best-F1 scores for catalytic residues predicted by SCREEN. We compare SCREEN with results generated by a sequence-based baseline model (CNN encoder). The native catalytic residues are in green in a zoomed figure. The correctly identified catalytic residues by SCREEN and a sequence-based baseline model (CNN encoder) are marked in red, while misidentified residues are in gray.

Using SCREEN, we also measured Best-F1 score and AUPR values for specific enzyme types (Supplementary Fig. 8). Particularly, for hydrolases, which represent the largest portion of the training dataset (313 of 1055; 29.7%), SCREEN obtained notable consistency across the five test datasets, with Best-F1 scores ranging from 0.63 to 0.78, AUPR values ranging from 0.56 to 0.68. For the isomerase datat, the predictive performance was particularly high, with the Best-F1 score exceeding 0.74 and AUPR surpassing 0.80, even though these enzymes represent only a small portion of the training set (87 of 1055; 8.2%).

### The use of enzyme structure information in SCREEN markedly improves the catalytic residue prediction

The catalytic residues typically tend to form cohesive clusters within the three-dimensional enzyme structures. Thus, we systematically investigated the spatial distribution of residues in enzyme structures by measuring the Euclidian distances to the nearest catalytic residue for both catalytic and non-catalytic residues in individual protein sequences. Fig. 2b shows there was a clear difference in the distribution of Euclidian distances, with the catalytic residues peaking at ∼ 6Å, and most non-catalytic residues exceeding 15Å, supporting that catalytic residues form cohesive active enzymatic sites. We investigated whether SCREEN could reconstruct the same spatial distributions for enzyme catalytic residues. A performance assessment of SCREEN using five test datasets (Fig. 2c) showed that the majority of catalytic residues predicted grouped together in the structure and that the distance values were consistent among the datasets, with median values being ∼ 6Å. These results agree with findings presented in Fig. 2b, implying SCREEN accurately captures the spatial distribution of predicted catalytic residues.

These findings suggest the use of structure information in SCREEN likely results in predictive performance improvements. We further investigated whether SCREEN could improve results compared with a ‘baseline model’ that excludes structure-based inputs and replaces the graph convolutional neural network (GCN) with a sequence-based convolutional neural network (CNN). The comparison of SCREEN with the baseline using five test datasets employing the Best-F1 score, AUPR and AUC metrics (Fig. 2d) showed that the use of graph network led to a marked improvement in predictive performance.

To assess predictions, we selected enzymes from the EF superfamily dataset ranked around the 10%, 50%, and 90% percentiles based on the Best-F1 score. We plotted catalytic residues predicted by the structure-based SCREEN and the sequence-based baseline model against ground-truth catalytic residues within enzyme structures. In the top 10%, for carboxylic ester hydrolase (PDB ID:1LE6), SCREEN accurately predicted all catalytic residues, whereas the sequence-based baseline introduced three false positives (Fig. 2e). In the median ranking, for casein kinase-1 (PDB ID:1CSN), SCREEN successfully identified Asp-131, Lys-133 and Thr-181 as key residues, but the baseline model failed to identify Thr-181 (Fig. 2f). In the 90% ranking, for dihydropteroate synthase (PDB ID:1AD1), SCREEN identified Arg-239 and gave one false positive, whereas the baseline misidentified two non-essential residues (Fig. 2g). These findings indicate that SCREEN has a superior performance compared with the sequence-based baseline model. This supports our design and, in particular, the use of the graph network and enzyme structure as a key input. We also investigated various graph convolution types, including the extensively employed Graph Convolutional Layer (GCN), Graph Attention (GAT) and Graph Isomorphism Network (GIN), but none of the models outperformed another using the same test sets and metrics (Supplementary Fig. 6)

### SCREEN accurately predicts catalytic residues using structure models

Although we showed SCREEN’s predictive performance benefits from the use of enzyme structure data, this information is often missing for many proteins/enzymes. Recent advances in protein structure prediction, like the AlphaFold algorithm (*27-29*), make it possible to accurately predict protein structure from sequences and to use such structure models as the input to SCREEN. We evaluated whether the use of predicted structures rather than experimentally determined structures would alter SCREEN’s predictive performance. We generated *C*_*α*_-*C*_*α*_ contact maps for enzymes based on experimental protein structures sourced from PDB, as well as putative structures from AlphaFold. We tested SCREEN’s performance by using each of these two sets of contact maps, comparing it against a sequence-based baseline model devoid of structural information. Fig. 3a showed that SCREEN consistently benefits from utilizing putative enzyme structures (with Best-F1 = 0.702, AUPR = 0.644, and AUC = 0.985) compared with sequence data alone (with Best-F1 = 0.691, AUPR = 0.624, and AUC = 0.973).

**Fig. 3.**
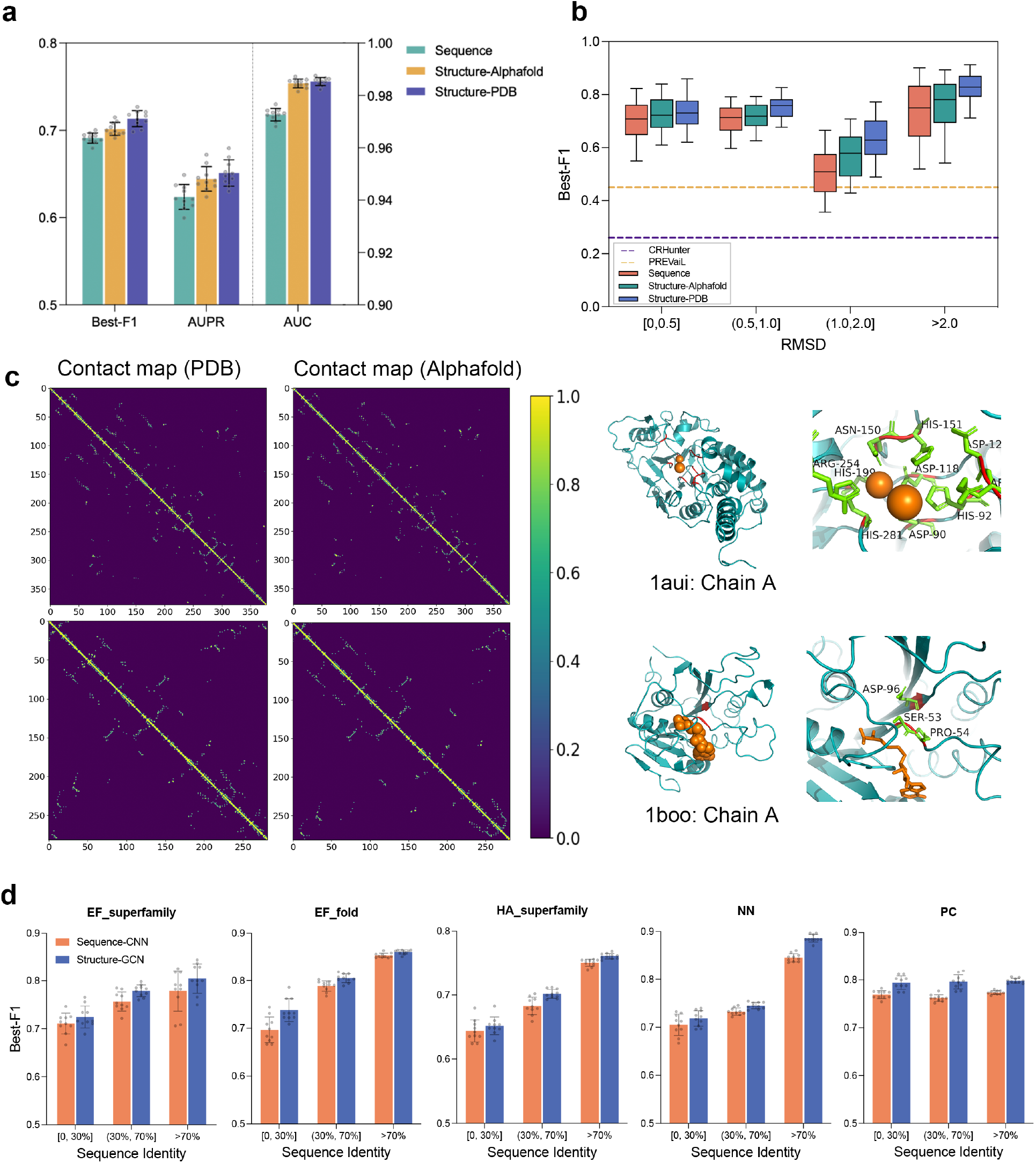
Analysis of predictive performance when using putative enzyme structure and low similarity test proteins. **(A)**, Comparison of results produced by the SCREEN using the native structure, structure predicted from sequence with AlphaFold, and sequence-based baseline predictor (CNN encoder); the error bars represent standard deviation of the mean based on 10 independent runs. **(B)** Comparison of results produced by the SCREEN using the native structure, AlphaFold-predicted structure, and sequence-based baseline predictor in the context of the quality of the AlphaFold-predicted structure (x-axis). The boxplots show distributions of the per-enzyme Best-F1 scores. The dashed horizontal lines represent the Best-F1 scores generated by the sequence-based CRHunter (blue line) and the recently published PREvaIL (yellow line). **(C)** Examples of contact maps by ground-truth of native enzyme structures (PDB) and AlphaFold-predicted enzyme structures, and corresponding catalytic residues identified by SCREEN for enzymes 1aui-A and 1boo-A. **(D)** Evaluation SCREEN predictions for enzymes that share varying levels of sequence identify with the training proteins using AUPRC, Best-F1 scores on the five test datasets.

To further assess SCREEN’s denoising power on predicted structure error, we evaluated the model performance employing AlphaFold-predicted structures with varying quality. Specifically, we quantified the quality of predicted structures employing root-mean-square deviation (RMSD) metric as compared with experimental solved structures (RMSD=0). Fig. 3b revealed that SCREEN’s performance was better employing AlphaFold-derived structures than using sequence data alone. SCREEN also out-performed currently-employed tools CRHunter and PREVAIL across the entire RMSD range, achieving the Best-F1 score of > 0.6 using AlphaFold-predicted structures, contrasting average Best-F1 scores of 0.45 and 0.26 for CRHunter and PREvaIL, respectively.

SCREEN performed relatively well using predicted structures (Fig. 3a, 3b), irrespective of the quality of predictions via AlphaFold. Here, we used two examples to illustrate SCREEN’s ability to accurately identify catalytic residues even in relatively low-quality predicted structures (Fig. 3c). For the human calcineurin heterodimer (PDB ID: 1AUI, Chain A) (*30*), where the AlphaFold-predicted structure had an RMSD score of up to 6.76, SCREEN successfully identified all 10 catalytic residues. Similarly, SCREEN accurately identified catalytic residues (Ser-53, Pro-54 and Asp-96) in the PVUII DNA methyltransferase (PDB ID: 1BOO, Chain A) (*31*), for which the structure predicted had an RMSD score of 4.75. Taken together, these results suggested that SCREEN can accurately predict catalytic residues from AlphaFold-predicted enzyme structures, which might be attributed to the robustness of the input features and how they are represented in the graph network model.

### SCREEN predicts catalytic residues in “previously-unseen” enzymes

We investigated SCREEN’s ability to generate accurate predictions for “previously unseen” enzymes. To this end, we categorized enzymes in the five test sets into three distinct groups based on their sequence identities, namely ≤ 30% (low), 30 to 70% (moderate) and > 70% (high), to enzymes in the training dataset. Fig. 3d showed SCREEN’s Best-F1 scores across three sequence identity ranges, i.e., ≤ 30% (low), 30 to 70% (moderate), and > 70% (high), for each of the five test datasets. We showed that SCREEN consistently outperformed the sequence-based baseline model for each of the five datasets and the three identity ranges. Importantly, SCREEN’s predictions were accurate also for enzymes with limited sequence identities to those in the training datasets, achieving the Best-F1 scores of 0.725, 0.738, 0.652, 0.718, and 0.794. These predictions were significantly better than those achieved using the existing tools, such as CRHunter (Best-F1 values of 0.328, 0.315, 0.450, 0.365 and 0.140, respectively) and PREVAIL (Best-F1 of 0.264, 0.261, 0.263, 0.264 and 0.263, respectively).

### Model training for enzyme classes improves the prediction of catalytic residues

We investigated whether the training of the graph network model with distinct enzyme function implications would improve predictive performance, considering that the small subset of catalytic residues contributes to the intricate functions of enzymes. We used enzyme class information (first-level EC numbers) to refine enzyme representation through a contrastive learning framework during the training process. This led to a separation of latent feature spaces in SCREEN’s deep network model for different types of enzymes. We displayed these latent feature spaces among different enzyme classes employing t-distributed stochastic neighbor embedding (t-SNE) (*32*). We showed that predictive performance (metrics: F1, AUPR and AUC) of SCREEN was enhanced compared to when enzyme function was not incorporated for each of the data sets (EF fold, HA superfamily, EF superfamily, NN and PC) (Fig. 4a and Supplementary Fig. 11) and that SCREEN was able to group enzymes with similar functions together and separating enzymes with distinct functions (Fig. 4c). Taken together, these findings indicate that the enzyme function incorporation through contrastive learning during the training process improves predictive performance as SCREEN can differentiate catalytic from non-catalytic residues for different distinct types of enzymes.

**Fig. 4.**
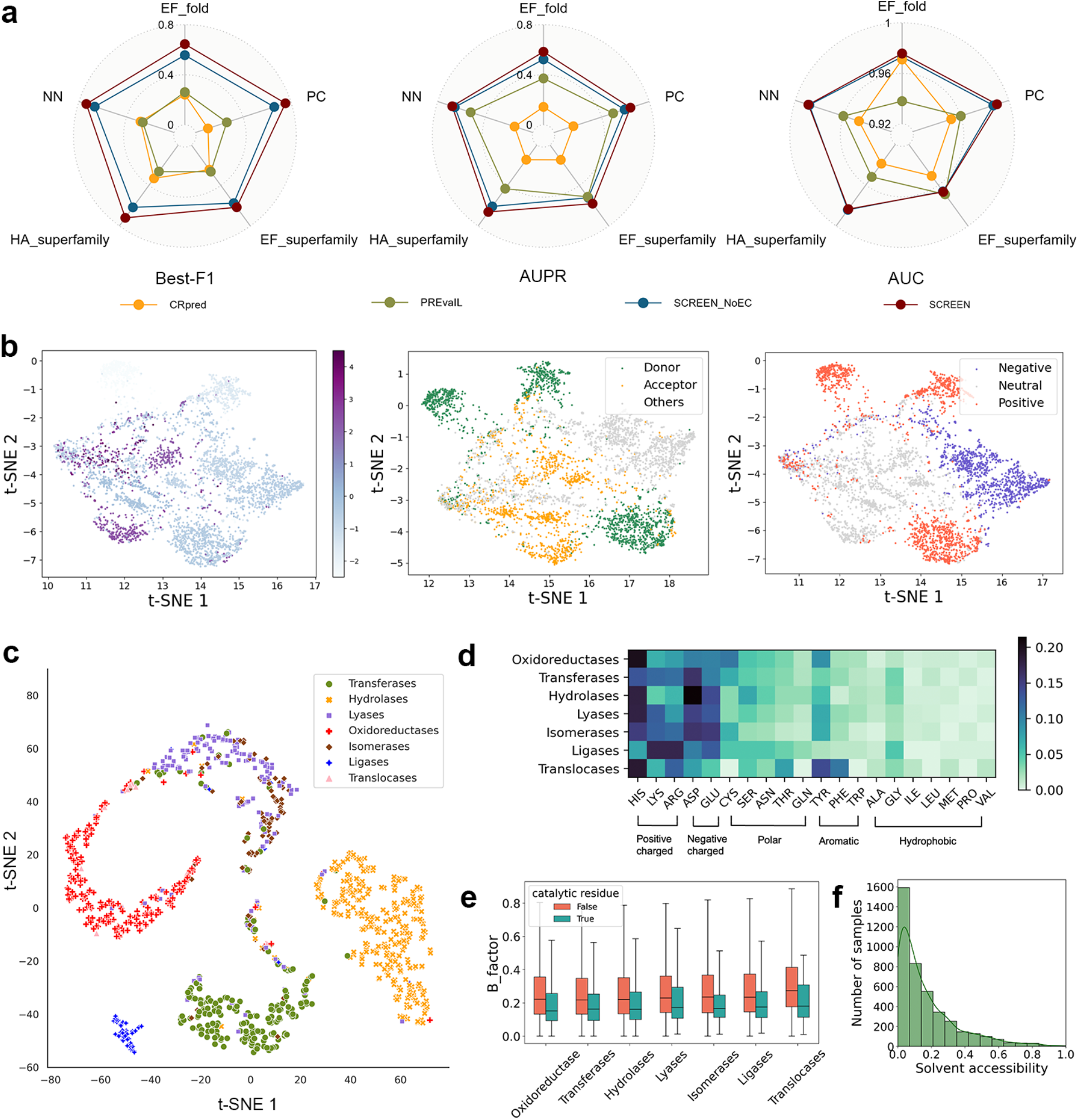
Analysis of putative catalytic residues generated by SCREEN. (**A**) Comparison of the SCREEN models with its variant SCREEN_NoEC that does not incorporate enzyme function through contrastive learning using Best-F1, AUC and AUPRC metrics on the five test datasets. As a point of reference, this figure also shows results from the sequence-based CRpred and the recent structure-based PREvaIL. (**B**) The t-SNE based visualization of latent feature spaces in the SCREEN model for residue characteristics, such as hydrophobicity (left), hydrogen bond types (medium) and charges (right). The “H-bond acceptor” denotes residues exclusively serving as hydrogen bond acceptors without containing H-bond donor atoms. (**C**) The t-SNE based visualization of latent feature spaces in the SCREEN model for different color-coded enzyme classes. (**D-F**), Analysis of structural and biophysical characteristics, which include biophysical properties of amino acids (D), B-factor (E) and solvent accessibility (F) for the putative catalytic residues generated by SCREEN.

### SCREEN can capture selected features of catalytic residues

We analyzed catalytic residues predicted by SCREEN, in order to investigate whether they possess structural and biophysical characteristics expected for enzymes. To better understand the relevance of the features learned by SCREEN, we initially displayed the general chemical properties (including hydrophobicity, charge and hydrogen bonds), along with low-dimensional projections of residue-level representations (Fig. 4b). We observed that charged amino acids dominated in the catalytic residues predicted (Fig. 4d), consistent with previous findings showing that electrostatic filtering has a marked effect on enzyme substrate selection (*33, 34*). Moreover, catalytic residues were inferred to be more rigid (structurally) than non-catalytic residues (based on low *vs*. high B-factor values; see Fig. 4e), which accords with a previous study of native catalytic residues (*35*). Fewer hydrophobic residues were associated with catalytic residues (Fig. 4d), which is consistent with limited solvent accessibility (Fig. 4f) and suggests substrate avoidance in substrate–enzyme interactions (*34*). Collectively, these results show that SCREEN can capture key features that typify native catalytic residues in distinct classes of enzymes.

### Linking catalytic residues to enzyme function and structure

Based on these catalytic residues, we further analyzed the sequence-structure-function relationship of enzymes to gain deeper insights into enzymes’ catalytic mechanisms. We categorized enzymes according to their catalytic functions defined by third-level EC numbers and then by (complete) fourth-level EC numbers (which link to substrates). For enzyme clusters sharing the same catalytic function/mechanism, we assessed structural similarity by their TM-scores among cluster members (*36*), selected enzymes from individual clusters and mapped the catalytic residue predictions to respective three-dimensional structures (Fig. 5).

**Fig. 5.**
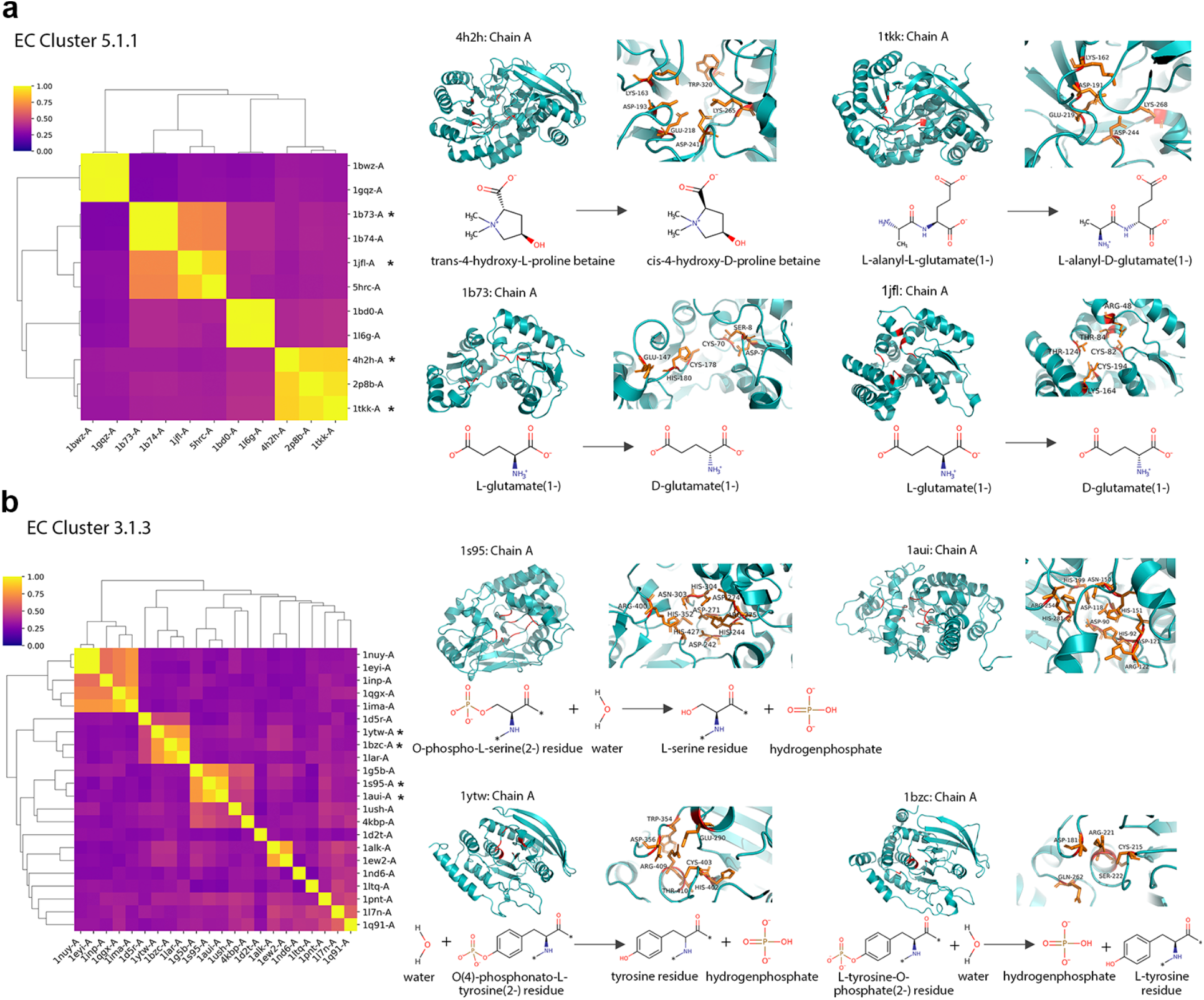
Diversity of enzyme structure and catalytic residues with the same catalytic mechanism. We examine enzymes that have the same catalytic mechanism: (A) Isomerases acting on amino acids and derivatives with EC number 5.1.1 and (B) phosphoric monoester hydrolases with EC number 3.1.3. We plot the TM-score as a measure of structural similarity as a heatmap, with larger numbers (more yellow) representing more similar structures. We also map the catalytic residues prediction by SCREEN onto the structures as well as enzyme reactions on the right.

The results indicated that same catalytic motif may represent distinct enzymes that serve different functions linked to diverse ligands or substrates. Fig. 5a shows both 4-hydroxyproline betaine 2-epimerase (PDB ID: 4h2h, Chain A) and L-Ala-D/L-Glu epimerase (PDB ID: 1TKK, Chain A) belong to the same superfamily and share a common catalytic motif (KDEDK) (*37, 38*), but they differ in their substrate specificity: 4-hydroxyproline betaine 2-epimerase facilitates the 2-epimerization of trans-4-hydroxy-L-proline betaine (tHyp-B) to cis-4-hydroxy-D-proline betaine (cHyp-B), whereas L-Ala-D/L-Glu epimerase catalyzes the reversible epimerization of L-Ala-D-Glu to L-Ala-L-Glu.

Enzymes in a particular family can have similar structures and functions, despite undergoing sequence divergence through evolution. A compelling illustration emerges when we studied two related enzymes, the glutamate racemase (PDB ID: 1b73, Chain A) (*39*) and aspartate racemase (PDB ID: 1jfl, Chain A) (*40*) (Fig. 5a). Despite enabling similar reactions via the same mechanism, their catalytic residues are significantly different but have analogous tertiary structures. Another example relates to protein-tyrosine-phosphatase non-receptor class (PDB ID: 1ytw, Chain A) (*41*) and protein-tyrosine-phosphatase non-receptor type 1 (PDB ID: 1bzc, Chain A) (*42*) (Fig. 5b). Here, although both enzymes are tyrosine phosphatases and catalyze the same reaction to remove phosphoryl groups from tyrosine residues in proteins, their respective catalytic residues are distinctly different.

The shared structural arrangements of catalytic residues can be associated with functional similarity. Fig. 5b shows phosphatase 5 (PDB ID: 1S95, Chain A) and phosphatase 2B (PDB ID: 1aui, Chain A) exhibite significant structural similarity and share catalytic residues (motif: DHDDRNHHRH) pertaining to serine/threonine phosphatase function(s), characterized by executing a “nucleophilic assault” on the phosphorus atom within a phosphorylated serine or threonine residue (*30, 43*).

Although enzymes catalyzing the same reactions often exhibit marked sequence and/or structural similarity, exceptions exist where structurally dissimilar enzymes enable similar reactions via the same mechanism. This is expected since enzymes facilitate numerous reactions using a finite set of building blocks in their residues, resulting in multiple enzymes inevitably sharing components of their catalytic mechanisms. Here, we showed that non-homologous proteins, protein phosphatase 5 (PDB ID: 1S95, Chain A) and dual-specificity phosphatase (PDB ID: 1d5r, Chain A) (*47*) employ distinct structural motifs to execute the same reaction that dephosphorylates a phosphoprotein substrate (Fig. 5b and Supplementary Fig. 12).

### Associating mutations with the SCREEN-predicted catalytic pockets

Here, we studied the patterns of mutations in the context of their proximity to the catalytic pockets predicted by SCREEN. We collected the MAVE data, probing mutation effect(s) on the functions of four different enzymes, namely PTEN tumor suppressor (PDB ID: 1d5r, Chain A) (*44*), human cytochrome P450 CYP2C9 (PDB ID: 1og5, Chain A) (*45*), NUDT15 (PDB ID: 5lpg, Chain A) (*46*), and *Escherichia coli* TEM1 beta-lactamase (PDB ID: 1btl, Chain A) (*47*), encompassing a total of 15,665 variants across 1,343 residues.

To function effectively, enzymes must be present at sufficiently high levels and have suitable catalytic residues in the active sites (*48*); however, mutations can affect both of these aspects, potentially resulting in impaired enzymatic function. We used the MAVE data (*49*) for two residue classes: (1) wild type-like (WTL) residues that exhibit high functional tolerance to mutations, whose most missense mutations do not adversely impact enzyme function; and (2) functional loss (FL) residues that are prone to mutations that either decrease abundance (e.g., unstable structures) and/or impair function, leading to diminished enzyme activity.

We systematically analyzed key characteristics of mutations in the context of the predicted catalytic residues. Specifically, we applied an additional tree-structured model to SCREEN (Fig. 6a). The Euclidean distance values to the closest catalytic residue predicted by SCREEN combined with solvent accessibility, which, as expected, were inferred to vary according to residue type (WTL or FL), allowing to differentiate among different mutation groups. Fig. 6b illustrates the catalytic residues predicted by SCREEN along with residue mutation type predictions. We performed five-fold cross-validation on all MAVE data; our results revealed an average accuracy of 0.70 and 0.84, and precision of 0.58 and 0.88 on the validation data and the entire dataset, respectively (Fig. 6c). We found distinct spatial distribution patterns for the WTL and FL residues based on their Euclidean distances from the putative catalytic residues (Fig. 6d). Our result aligns well with experimental data showing that FL residues are relatively close to the catalytic site, while WTL residues are distributed throughout the structure (Supplementary Fig. 13). This result indicates that SCREEN can be useful to establish the impact of mutations on enzyme structure and function and provides a tool to guide the identification of disease-associated mutations in enzymes.

**Fig. 6.**
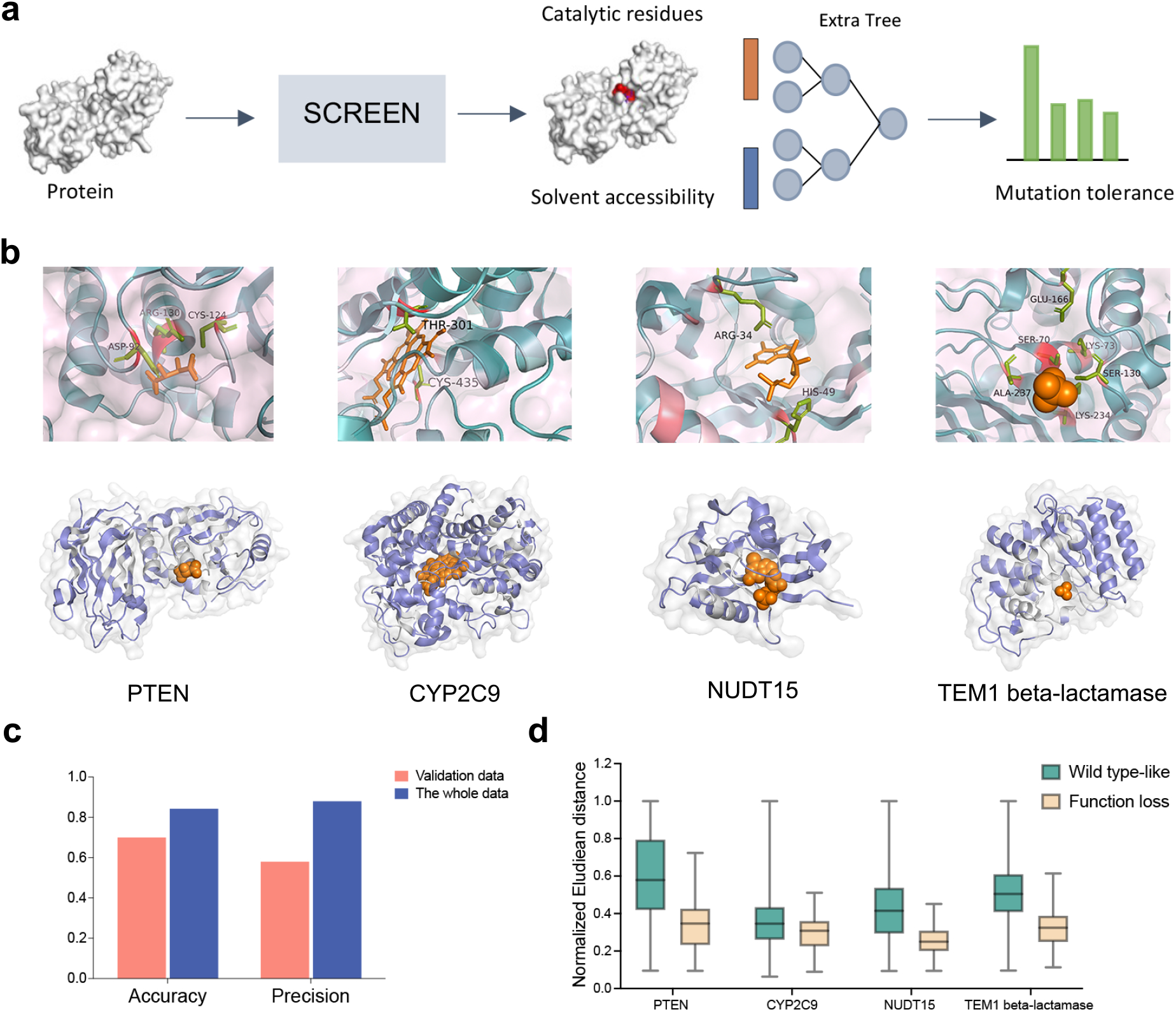
Assessing the tolerance of catalytic residues to mutations. (**A**) Architecture of the tree-structured model for characterizing mutated residues based on catalytic residue predictions by SCREEN. **(B)** SCREEN identified catalytic residues of four different enzymes: PTEN tumor suppressor (PDB ID: 1d5r, Chain A), human cytochrome P450 CYP2C9 (PDB ID: 1og5, Chain A), NUDT15 (PDB ID: 5lpg, Chain A), and *Escherichia coli* TEM1 beta-lactamase (PDB ID: 1btl, Chain A) (top). Residues within enzyme structures are colored according to their predicted mutant class: blue corresponds to the Wild Type-like (WTL) residues, while gray to the Functional Loss (FL) residues (bottom). **(C)** Quality of distance-based classification of residues with different mutation classes, measured by accuracy and precision on both the validation and entire datasets. **(D)** Distribution of the Euclidean distances for residues of different mutation classes.

## Discussion

Enzymes can catalyze a broad set of chemical reactions using a limited set of catalytic residues (*50*). Identifying these residues allows us to understand how existing enzymes function at the molecular level and to design new ones. In this work, we hypothesize that the structural organization of catalytic residues in spatial space, along with their generally high evolutionary conservation, collectively contributes to catalytic residue identification. To this end, we conceptualized, designed, and assessed SCREEN, a structure-based graph network that uses functional priors through contrastive learning and combines structure-, sequence-, and evolutionary profile-based representations to accurately predict catalytic residues in enzymes.

Comparative empirical assessments using five commonly-utilized test datasets and seven currently-available (published) predictors revealed that SCREEN (i) accurately predicts catalytic residues in known and computationally-modeled enzymes; (ii) outperforms current tools; and (iii) generalizes well to enzymes that have limited similarity to enzymes used to train the model, suggesting that SCREEN is applicable to currently-unknown enzymes. After demonstrating that SCREEN could infer key structural and biophysical features, including amino acid charge, solvent accessibility and structural rigidity, of predicted and known catalytic residues, we undertook sequence-structure-function analyses to link catalytic residues to enzyme structure and function. Our analyses revealed that while enzymes that catalyze identical reactions often display significant sequence and/or structural similarity, exceptions arise wherein dissimilar sequences and/or structures can catalyze reactions via the same mechanism. In addition, using multiplexed enzyme mutation data, we showed that SCREEN could infer the tolerance of individual catalytic residues to mutations and, thus, predict which mutations in catalytic residues likely lead to the functional loss of an enzyme. Taken together, SCREEN should provide a useful tool for the reliable prediction of catalytic residues to support studies of known and unknown enzyme groups/classes as well as enable *in silico* investigations of diseases linked to mutations.

## Materials and Methods

### Training and test datasets

We curated a dataset comprising 1055 enzymes with annotated catalytic residues, which we used to train and optimize our predictive model. First, we collected data from M-CSA database, which contains catalytic residue annotations and details about enzymic reaction mechanisms (*51*). We combined the EF family dataset that includes catalytic residue annotations for enzyems from different SCOP families (*20*). We filtered the combined dataset by clustering the proteins with the CD-HIT software at 40% sequence identity (*52*), and picking one protein from each cluster. This prevented overfitting the model into larger clusters of similar enzymes. Next, we collected enzyme three-dimensional structures from the Protein Data Bank (PDB) (*9*). We obtained the Enzyme Commission (EC) numbers (*53*) using the SIFTS program (*54*). We used the first-level EC numbers due to the relatively sparse/incomplete nature of these data at the lower levels (*55*). We shuffled the dataset randomly and then divided it into training (90%) and validation (10%) subsets.

We acquired five widely-used test datasets to conduct a comparative evaluation of our model against existing tools (*16, 18, 20, 22*). These testsets included the EF-fold dataset, EF-superfamily dataset and HA superfamily dataset, representing enzymes from every SCOP fold and superfamily respectively. We also collected the PC test dataset (*18*), which was originally obtained from Catalytic Residue Dataset (CATRES) and represents proteins of the PIRSF protein groups (*56*). Finally, we obtained the NN test dataset (*16*), which comprises enzymes of six main classes. Importantly, we excluded enzymes from these five test datasets from our training/validation dataset. Supplementary Table 1 summarizes the training and the five test datasets.

### Overview of the SCREEN model

The SCREEN model utilizes enzyme structures to accurately predict catalytic residues by leveraging a deep neural network and multiple heterogeneous inputs that include atomic structures, sequence embeddings that are based on a modern language model, and evolutionary information (e.g., conservation) (Fig. 1). The innovations of this model are: First, we used a geometrically invariant representation of the enzyme structure and sequence that captures three heterogeneous inputs (Fig. 1b). Second, we applied graph convolutional neural network (GCN)-based encoder as the predictive model (Fig. 1c). This network architecture is suited to the structure-derived data input by computation and the propagation of embeddings among residues which are spatially proximal in the structure. Third, we employed contrastive learning to utilize enzyme function information, which we combined with a dynamic strategy to train the SCREEN model.

### Graph-based representation of protein structure

We represented the input enzyme structure composed of n residues ***Enz***= {**r**_**1**_, **r**_**1**_,…, **r**_**n**_} as an attributed graph encoded with evolutionary context, sequence embeddings and relevant structural characteristics, such as solvent accessibility and B-factors. Specifically, the graph representation ***G*** = (***V, A***,***X***) consists of three-dimensional enzyme structure by taking the residue set ***V*⊆*Enz*** as graph nodes, the adjacency matrix ***A*** with ***n*** × ***n*** size that quantifies connectivity of nodes/residues, and the feature matrix ***X* ∈ *ℛ***^***n***×**θ**^.

The feature matrix ***X***= (***X***_***L***_,***X***_***G***_,***X***_***A***_) covers the evolutionary, sequence and structural features. The ***X***_***L***_ descriptor quantifies evolutionary conservation by utilizing two complementary tools: PSI-BLAST, which is a heuristic algorithm that relies on the dynamic programming (*23*), and HMMER that is based on the hidden Markov model (HMM) (*24*). We run PSI-BLAST on the NCBI’s non-redundant (nr) database, with three iterations and the E-value threshold of < **10**^**−3**^. We normalize the output position-specific scoring matrix (PSSM) of size ***n*** × **20** with the sigmoid function: 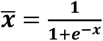. We use HMMER with the uniclust30 database (*57*) and default parameters to generate the ***n*** × **30** HMM matrix that we normalize to the [0, 1] range (*24*). The ***X***_***G***_ descriptor captures sequence information computed by ProtT5 model, a deep learning language model that was pre-trained on 390 billion amino acids (*58*). The enzyme sequence is encoded into residue-level feature embeddings denoted as 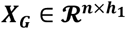, where **h**_**1**_ defaults to 1024. These vectors encapsulate information about individual residues that are adjacent in the sequence, and broader protein-level information. Lastly, the ***X***_***A***_ descriptor encompasses several key properties that are derived from the atomic-level data: atom types and atomic mass when excluding hydrogen atoms, ***B***-factor, residue side-chain presence, the count of bonded hydrogen atoms, ring membership, van der Waals radius, and solvent accessibility. Given that residues might have different numbers of atoms, we compute the average values across all atoms, resulting in atomic descriptor 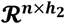, with **h**_**2**_ = 14.

### Predictive model

We designed the graph convolutional neural network (GCN) with three convolutional layers (*59*) to facilitate the propagation of feature embeddings for residues that share spatial proximity.

For a given graph defined by the adjacency matrix ***A* ∈ {0, 1}**^***n***×**n**^ and the feature matrix ***X***= (***X***_***L***_,***X***_***G***_,***X***_***A***_), our model produces residue-level representations 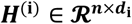 where ***d***_**i**_ represents the embedding dimension for the ***i***th convolutional layer.

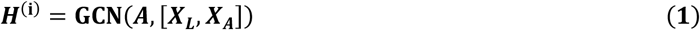

We refine residue representations through the process of neighbor aggregations as follows:

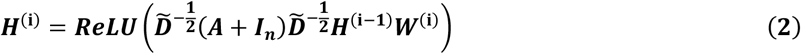

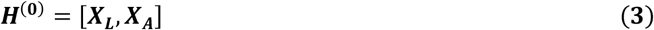

where ***I***_***n***_ **∈ *ℛ***^***n***×**n**^is the identity matrix, 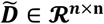 is the diagonal degree matrix with entries 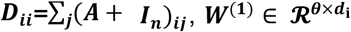 is the trainable weight matrix for the ***i***th convolutional layer, ReLU denotes the Rectified Linear Unit activation function, and [] denotes the concatenation operation. The above architecture generates graph representation ***X***_***E***_ **∈ *ℛ***^***n***×**d**^, where d =512, which we combine using multilayer perception (MLP) network as follows:

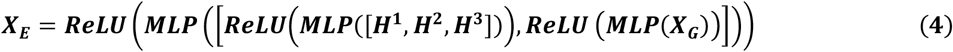

We employ three fully connected layers in the MLP network to reduce the feature space to the final output vector **Y ∈ R**^***n***×**2**^ that gives numeric propensities for putative catalytic residues.

### Contrastive learning

We used contrastive learning with Triplet Margin Loss to craft enzyme representations that improve the catalytic residue predictions. Using the graph representation ***X***_***E***_ **∈ *ℛ***^***n***×**d**^, we employed average aggregation across residues, generating a fixed-sized sequence representation vector 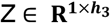, with **h**_**3**_ set to 1024. During training in each epoch, we iteratively refined every sequence representation vector and computed enzyme class cluster centres. When training with a query enzyme ***z***_***a***_, we selected the enzyme cluster centre embedding from the same enzyme class as the positive sample ***z***_***p***_, and randomly sampled another cluster centre from a different enzyme class as the negative sample ***z***_***n***_, which resulted in the following Triplet Margin Loss function:

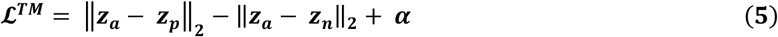

where we set the margin α to the default value of 1. This loss function minimizes the Euclidean distance between enzyme representations belonging to the same main enzyme class while maximizing the distance between those form different main enzyme classes. We implemented a dynamic training strategy, where we performed contrastive learning for enzyme classification during early training epochs, and gradually shifted towards the default training that converges to produce accurate propensities for catalytic residues.

### MAVE data analyses

We gathered MAVE measurements for four enzymes, which provided insight into the impact of a broad collection of substitutions on both enzyme function (*6, 60*). We categorized the missense variants of PTEN into two main groups: functional or inactive, regardless of their effect on abundance. The classification thresholds for the scores generated by each MAVE were guided by an established methodology (*61*). Specifically, we used a minimal number of Gaussians (three) to ensure a reliable fit to the variant score distributions, and the intersection point between the first and last Gaussian served as the classification cut-off. Adopting this binary classification approach allowed us to categorize variants into two classes: (1) Wild Type-Like (WTL): variants characterized by high activity; (2) Functional Loss (FL): variants assigned that exhibit low activity.

## Data availability

All raw and benchmark data were obtained from the following databases: Protein Data Bank (https://www.rcsb.org), UniProt (https://www.uniprot.org), SIFTS (http://pdbe.org/sifts/), and M-CSA (https://www.ebi.ac.uk/thornton-srv/m-csa/). All source data needed to evaluate the conclusion s in the paper can be found at https://huggingface.co/datasets/Biocollab/SCREEN/tree/main.

## Code availability

All source code is available at https://github.com/BioColLab/SCREEN.

## Acknowledgments

Financial support to RBG and JS from the Australian Research Council (ARC LP220200614) is gratefully acknowledged.

